# Establishing comprehensive quaternary structural proteomes from genome sequence

**DOI:** 10.1101/2024.04.24.590993

**Authors:** Edward Alexander Catoiu, Nathan Mih, Maxwell Lu, Bernhard Palsson

**Affiliations:** Department of Bioengineering, University of California, San Diego, La Jolla, CA 92101; Omnicorp Inc. (Pilot AI), San Francisco, CA 94129; The Novo Nordisk Foundation (NNF) Center for Biosustainability, The Technical University of Denmark, Kongens Lyngby 2800, Denmark

## Abstract

A critical body of knowledge has developed through advances in protein microscopy, protein-fold modeling, structural biology software, availability of sequenced bacterial genomes, large-scale mutation databases, and genome-scale models. Based on these recent advances, we develop a computational framework that; i) identifies the oligomeric structural proteome encoded by an organism’s genome from available structural resources; ii) maps multi-strain alleleomic variation, resulting in the structural proteome for a species; and iii) calculates the 3D orientation of proteins across subcellular compartments with residue-level precision. Using the platform, we; iv) compute the quaternary *E. coli* K-12 MG1655 structural proteome; v) use a dataset of 12,000 mutations to build Random Forest classifiers that can predict the severity of mutations; and, in combination with a genome-scale model that computes proteome allocation, vi) obtain the spatial allocation of the *E. coli* proteome. Thus, in conjunction with relevant datasets and increasingly accurate computational models, we can now annotate quaternary structural proteomes, at genome-scale, to obtain a molecular-level understanding of whole-cell functions.

**Significance:** Advancements in experimental and computational methods have revealed the shapes of multi-subunit proteins. The absence of a unified platform that maps actionable datatypes onto these increasingly accurate structures creates a barrier to structural analyses, especially at the genome-scale. Here, we describe QSPACE, a computational annotation platform that evaluates existing resources to identify the best-available structure for each protein in a user’s query, maps the 3D location of actionable datatypes (*e.g.*, active sites, published mutations) onto the selected structures, and uses third-party APIs to determine the subcellular compartment of all amino acids of a protein. As proof-of-concept, we deployed QSPACE to generate the quaternary structural proteome of *E. coli* MG1655 and demonstrate two use-cases involving large-scale mutant analysis and genome-scale modelling.

## Introduction

The proteome of the cell is responsible for metabolite uptake and secretion, genetic information processing and replication, energy production, and all other processes required for maintaining cellular homeostasis. Before becoming a functional unit of this multi-scale system, a protein must fold properly into its native three-dimensional shape. This folding process is also multi-scale. A peptide sequence (primary sequence) associates locally to form small recognizable patterns (e.g., alpha-helices and beta-sheets). Often stabilized by disulfide bridges and physiochemical attraction, these secondary structures fold onto each other to form larger recognizable domains, resulting in the three-dimensional structure of the protein monomer (tertiary structure). These protein monomers often oligomerize and form multi-subunit enzymes (quaternary structures) that carry out the functions in the cell.

Structural biology—the study of protein shape and function—has advanced rapidly in recent years. For proteins that form large multi-subunit complexes and for those spanning the cell membrane, the three-dimensional shape was particularly difficult to study with classical crystallographic techniques. The development of cryogenic electron microscopy – a method that images thin slices of a protein frozen in its native state (much like a biopsy)—has drastically increased the speed and ease by which these previously unknowable protein structures can be resolved^1–5^. Concurrently, computational methods have also experienced increasing success in accurately predicting protein structures^6–10^. Most recently, deep learning algorithms^11–12^ (e.g., AlphaFold) have utilized multiple sequence alignments and incorporated biophysical knowledge about protein structure to predict the shape of proteins without homologous structures in the Protein Data Bank^13^. Even more promising, these algorithms can be “hacked” to predict the structures of oligomeric assemblies of protein complexes^14^. Recent benchmarking efforts have confirmed the accuracy of homodimer AF-multimer models^15^ and have subsequently been used for homo-oligomeric predictions^16^.

Although structural biology can offer molecular insights into a protein’s shape and function, mutations in key domains can change the enzymatic properties and modulate a protein’s function. Changes in protein function can be either beneficial (e.g., an increase in stability of the active form) or detrimental (e.g., a loss of substrate binding efficiency). Protein engineering employs a variety of techniques to find mutations that produce a desired phenotype. One such technique, multiplex automated genomic engineering (MAGE)^17^, can introduce many mutations with unknown effects at specific sites in the genome. Mutations that result in the desired phenotype can then be selected for. Laboratory evolution, an experimental approach involving the serial passage of a cell population in an increasingly stringent selection pressure, can speed up the evolutionary process and beneficial mutations can be identified by sequencing the endpoint strain^18–20^. The dramatic decrease in sequencing costs in the last ten years has allowed for many mutations identified by these experimental techniques to be collected in databases.

Concurrent with advancements in structural biology and the formation of large-scale mutation databases, systems biology—the study of systems-level cellular behavior— was driven by the development of genome-scale models (GEMs) of cellular metabolism that predict gene essentiality, growth phenotypes, and proteome allocation in a few organisms^21–26^. Software (e.g., ssbio^27^) was developed to map available structural information to these modeled proteomes. Structural systems biology—the study of structural biology at the systems-level—has incorporated protein structures into genome-scale models (GEM-PROs) to study protein-fold evolution and investigate structural differences between organisms^28–32^. Notwithstanding the incorporation of protein information at the monomer-level, the use of GEM-PROs is the most recent step towards building genome-scale models that reflect the physical nature of the cellular proteome.

Given the availability of high-quality protein structures and structural models that capture the shape of multi-subunit complexes, the deposition of mutation-phenotype information into large-scale databases, and the development of genome-scale models of cellular proteome allocation, the creation of a *genome annotation platform* with interoperability between structural, functional, mutational, and systems-level information is now possible.

In this study, we present the Quaternary Structural Proteome Atlas of a CEll (QSPACE) — a computational annotation platform that 1) utilizes state-of-the-art modeling software (e.g. Alphafold^11^ & Alphafold Multimer^14^) and the latest crystallographic depositions to identify a three-dimensional structural representation that accounts for the multi-subunit assembly of the cellular proteome; 2) calculates structural properties of the proteome; 3) provides a three-dimensional context to map functional information including enzymatic domains, binding sites, and protein interfaces; 4) draws mutational information from large-scale databases of laboratory-acquired mutations^20,33–35^ and of the wild-type natural sequence diversity (alleleome) of *E. coli*^36^; and 5) calculates the subcellular compartmentalization of the proteome with residue-level resolution.

The QSPACE platform allows users to rapidly interact with protein structural data for biological inquiries ranging from the single-protein to the genome-scale (GS). Using *E. coli* as an example, we present two separate genome-scale applications of the QSPACE platform to demonstrate its broad applicability. First, we exploit QSPACE’s superimposition of mutant datasets and annotated functional domains on the protein structure to calculate 100 residue-level features for over 12,000 published *E. coli* mutations in UniProt^37^, allowing us to build RF-classifiers capable of predicting the severity of amino acid substitutions. Second, we showcase how QSPACE’s subcellular compartmentalization of the protein structures advances genome-scale modelling efforts. By calculating the size (volume, and, when applicable, the cross-sectional area of membrane proteins) of *E. coli* protein structures and incorporating them into iJL1678b^26^—a genome-scale model that predicts the macromolecular expression (80%, by mass) of *E. coli* MG1655—we are able to predict the physical space (across multiple subcellular compartments) required by the computed proteome of *Escherichia coli* K-12 MG1655 at optimal growth rate. To our knowledge, this QSPACE/GEM-PRO is the most comprehensive whole-cell approach that captures the 3D nature of the *E. coli* structural proteome. As structural, mutational, and functional knowledge is discovered, and GEMs are developed with increasing specificity, QSPACE can provide a method to rapidly integrate all information related to the structural proteome for an increasing number of organisms. QSPACE can be deployed for any organism following the tutorial python notebook available at https://github.com/EdwardCatoiu/QSPACE/.

## Results

### Overview of the QSPACE platform

The Quaternary Structural Proteome Atlas of a Cell (QSPACE) is an annotation platform that compiles available structural data from the latest structural biology efforts to obtain a 3D representation of all codon positions in a genome – complete with residue-level biophysical, chemical, and mutational data (see Table S1 for details). The QSPACE of *E. coli* is presented as a CSV file in Dataset S1. The two user-defined inputs to the QSPACE platform (Fig. 1a) are i) a list of gene IDs and ii) a dictionary of protein complexes and the associated stoichiometric ratio of the genes that make up each complex (Dataset S2A). QSPACE automatically downloads all protein structures (and homology models) from RCSB-PDB^13^, ITASSER^8^, SWISS-MODEL^10^ & AlphaFold^12^ that correspond to any of the genes in the user-defined inputs. QSPACE then finds the 3D coordinate file (i.e. “structure”) that best reflects the user-defined (input #2) multi-subunit protein assembly (Fig. 1b, details in Fig. 2). When no available structures can accurately reflect the gene-stoichiometry of a protein complex, QSPACE will attempt to generate models for the protein structure using an external GoogleColab notebook running AlphaFold Multimer^14^ (v2.0 via ColabFold^38^).

**Figure 1:**
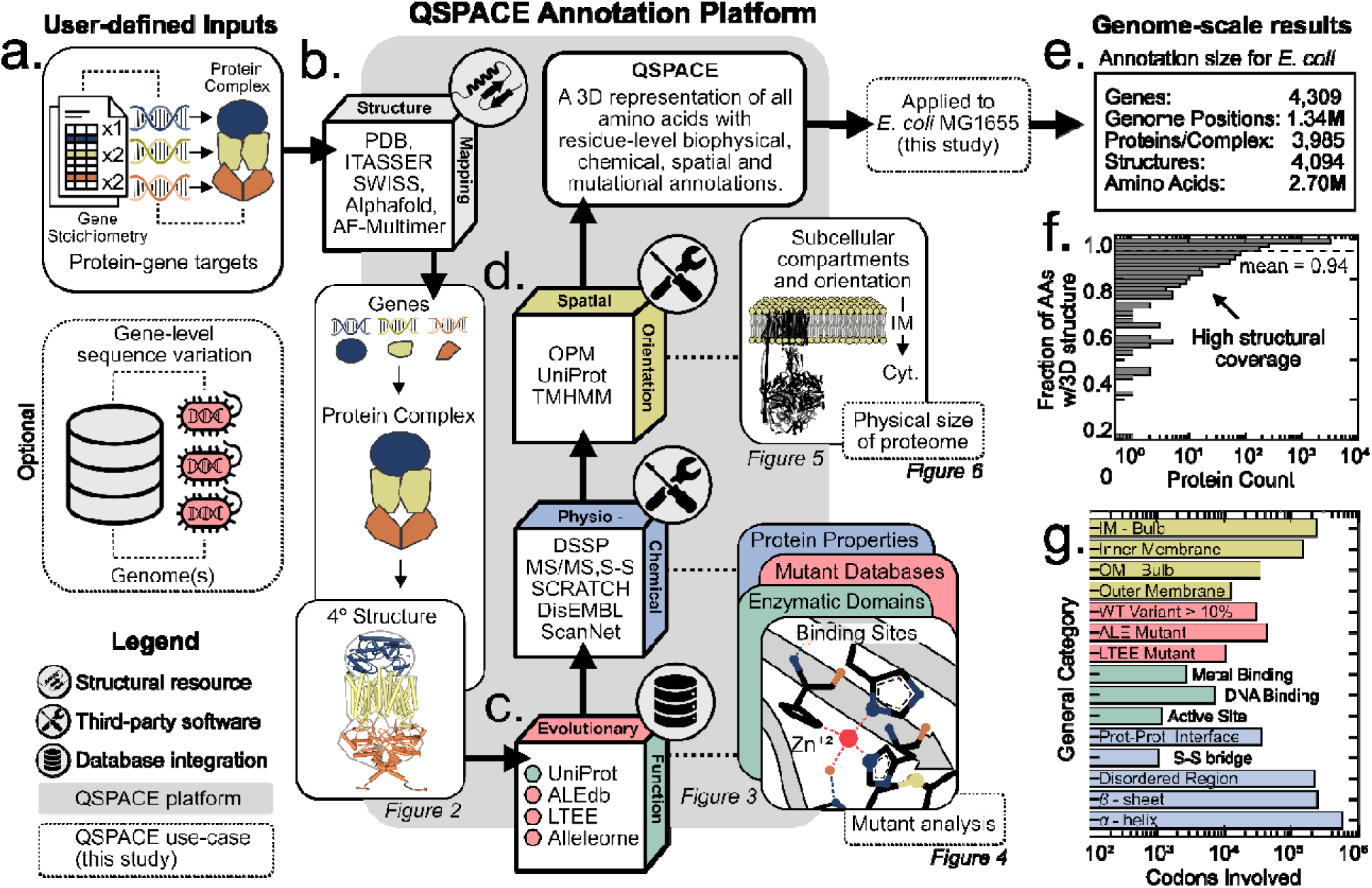
The Quaternary Structural Proteome Atlas of a CEll is a genome-scale annotation platform that was applied to the *E. coli* proteome. **(a)** The QSPACE platform requires two user-defined inputs: a list of gene(s) and dictionary of proteins and their associated gene-stoichiometric ratios. User-defined proteins can be oligomeric or monomeric. QSPACE accommodates residue-level sequence variation (the alleleome^36^ was used in this study). **(b)** QPACE identifies (or generates w/AF-multimer) the protein structure/model that best reflects the gene-stoichiometry of each user-defined protein complex (details in Figure 2a). The resulting structures **(c)** are analyzed using various software packages to calculate physicochemical properties and to identify evolutionary variable regions and functional domains (details in Figure 3a). **(d)** The structures are localized to their subcellular compartments and membrane-embedded structures are oriented across the membrane, resulting in a three-dimensional representation of **(e)** the structural proteome of an organism. **(f)** The amino acids of protein complexes are mapped to protein structures with varying levels of coverage (mean = 0.94) in *E. coli*. **(g)** Genome-scale counts of the unique codon positions belonging to various computed categories are shown.

**Figure 2:**
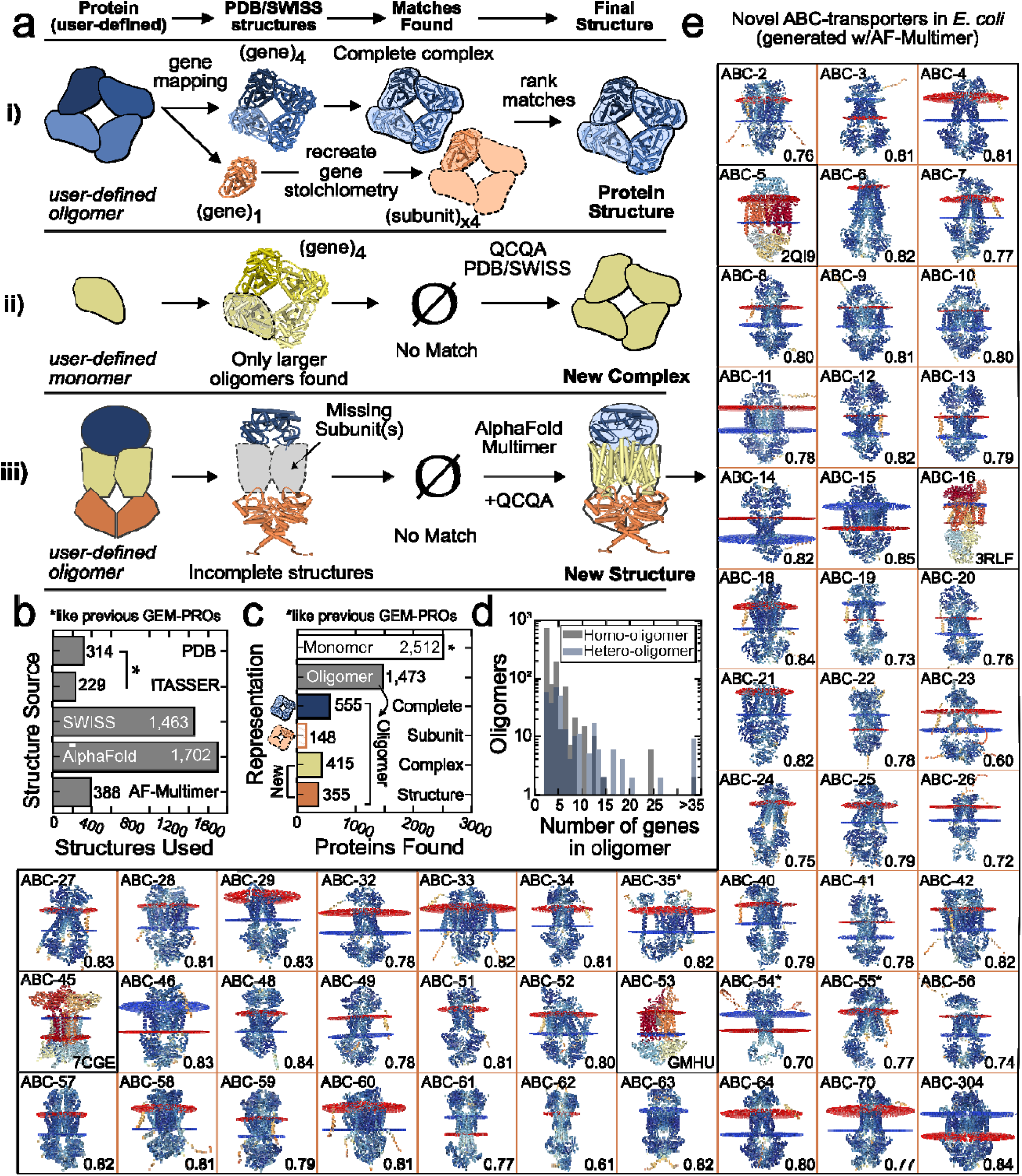
QSPACE yields 3D structures that reflect the oligomeric nature of multi-subunit proteins. (**a**) There are three ways that QSPACE’s protein-structure module finds the 3D structural representation for the user-defined gene-stoichiometry of a protein (target). (i) For each protein target (*left*), all structures that share genes with the protein are identified (*center left*) and combined (if applicable) to recreate the user-defined gene-stoichiometry (*center right*). If multiple matches are found, the structure that most accurately reflects the complete protein is selected (*right*). (ii) Sometimes, only higher-order structures of the protein-target are identified (*center left*). After QCQA of the relevant quality metrics associated with each structure (see Fig. S9, Methods 3.1), the highest confidence oligomeric structure is used to redefine the gene stoichiometry for the protein (‘new complex’, *right*). (iii) If identified structures are unable to recreate the protein in its entirety (‘missing subunits’, *center left)*, Alphafold Multimer is used to predict the structure (<2000 AAs) and the quality of the resulting oligomeric models are assessed (see Fig. S11, Dataset S9). (**b**) To achieve a multi-subunit representation of the *E. coli* proteome, structures or models from various sources are used. Unlike previous GEM-PROs, QSPACE accommodates monomeric and k-meric Alphafold models. (**c**) The protein-to-structure module yields 3D structures that represent the oligomeric nature of the *E. coli* proteome. This representation is further improved using structural data to correct existing protein-gene stoichiometry and with Alphafold Multimer to calculate novel oligomeric structures, (**d**) allowing for a truer accounting of higher-order oligomeric proteins. (**e**) The QSPACE platform generated 50 novel structures for ABC-transporters in *E. coli*. The pLDDT scores are used to color the AF-Multimer models. The associated AlphaFold Model Score (see Fig. S11, Dataset S3) is displayed in the bottom right. Asterisks are used to denote incomplete models resulting from incorrectly defined gene-stoichiometries or from protein size limitations of ColabFold. Existing PDB structures are shaded.

The thresholds used by QSPACE to assess the accuracy of selected protein structures are described in the text accompanying Figure 2 and in the methods section. All quality metrics related to protein structures for the *E. coli* QSPACE are provided in Dataset S3. By exploiting previously published repositories of protein structures, QSPACE reduces the threshold of interacting with genome-scale structural data to the order of days. Depending on user-preferences, QSPACE can function with all or some of the structural repositories used in this manuscript and can easily accommodate structural data from new sources as they become available.

Once the structure file representing the quaternary assembly of each protein is determined, multiple software packages and databases (see Table S1) are used to map physio-chemical, evolutionary, and functional information to the protein structures (Fig. 1c). The 3D overlay of multiple data types (details in Fig. 3) creates potential for many analysis tools (e.g., Fig. 4). The amino acids in each protein are then assigned to one of twelve subcellular compartments; and those representing the membrane fraction of the proteome are oriented across one of the *E. coli* membranes (Fig. 1d, details in Fig. 5). These structures can be integrated with genome-scale systems models to add a 3D understanding of the biophysical/spatial allocation of the proteome in a functioning cell (see Fig. 6). Users of QSPACE can bypass the mapping of any of these datasets if they are not relevant to their research, or not available for their organism.

**Figure 3:**
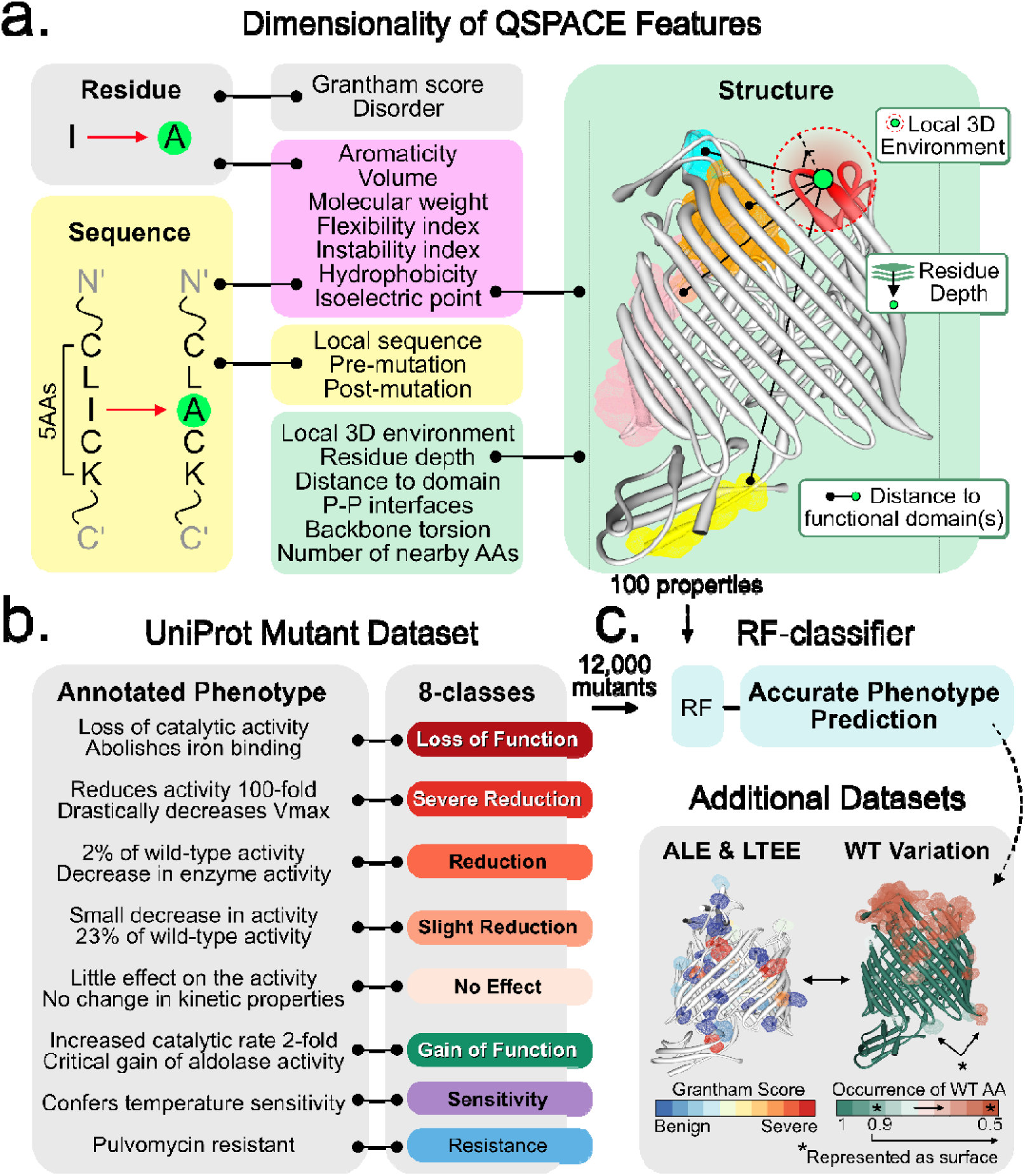
Multi-dimensional QSPACE features to predict mutant phenotypes. **(a)** QSPACE calculates 100 properties (residue-level, sequence-level, and structure-level) for all amino acids. (right) The mapping of the three-dimensional location of functional domains on the protein structure enables the calculation of a mutation’s proximity to important protein regions. **(b)** (left) UniProt mutations and their annotated phenotypes, are mapped to QSPACE and (right) can be classified into general categories that reflect mutant severity. **(c)** Annotated phenotypes and multi-dimensional properties for 12,000 UniProt mutations mapped to the *E. coli* QSPACE (Dataset S5) can be used to train RF-classifiers that can predict the effect of a mutation (see Fig. 4). We suggest the application of these classifiers to novel mutant datasets, (e.g. mutations in adaptive laboratory evolution experiments (ALE), the long-term evolution experiment (LTEE) and the alleleome are already mapped to the *E. coli* QSPACE in Dataset S1).

**Figure 4:**
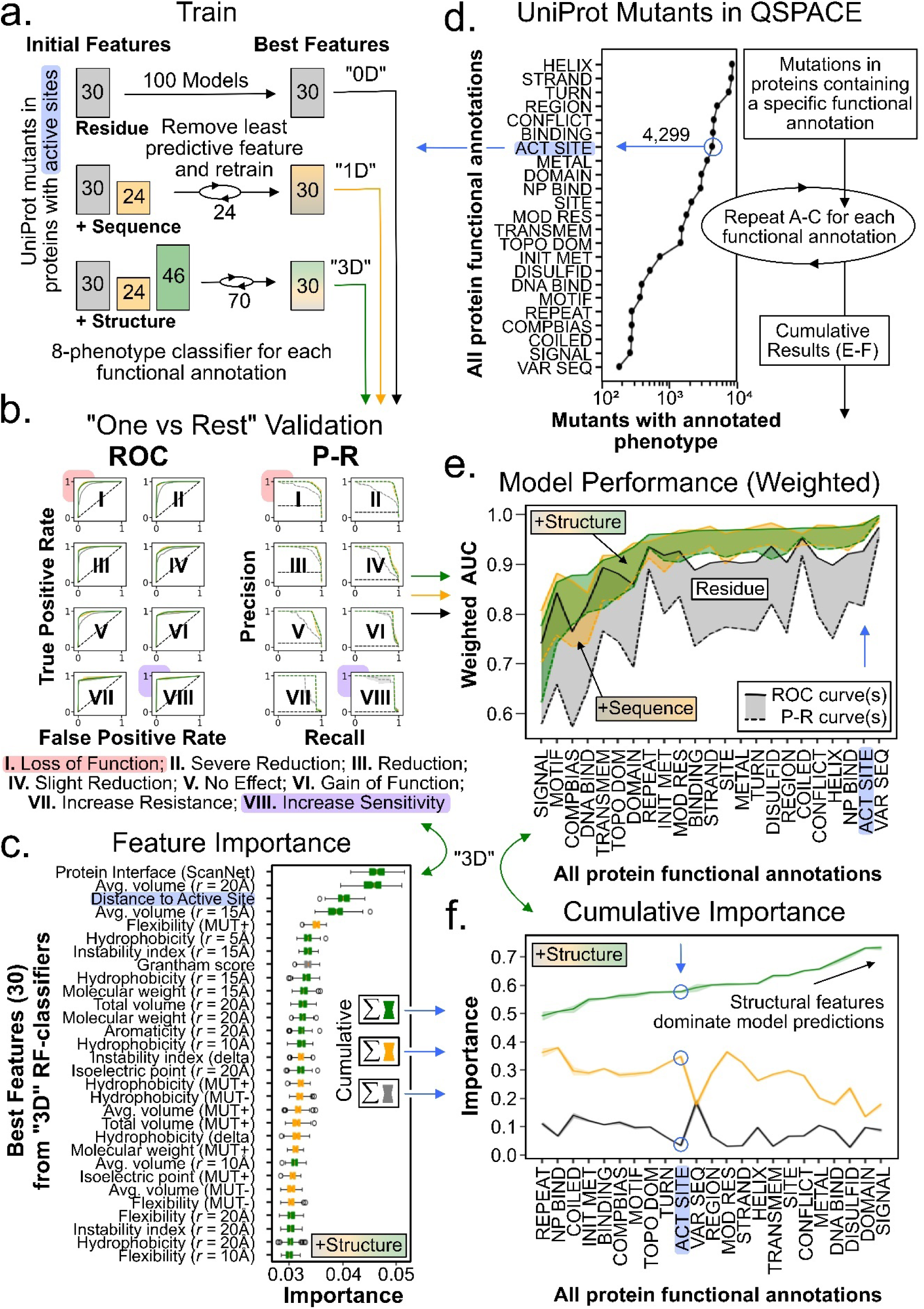
Random-forest prediction of mutant phenotypes in UniProtKB. **(a)** QSPACE finds 4,299 mutants with known phenotypes in proteins containing *active sites*. The mutational properties (‘features’) of the amino acid residue (grey), the 5 amino acid long sequence centered at the mutation (yellow), and the local 3D protein structure (green) are calculated (see Fig. 3). The numbers inside the boxes describe the number of features used. Random Forest Classifiers are trained on (top, “0D”) residue-level parameters; (middle, “1D”) residue and sequence-level parameters; and (bottom, “3D”) residue, sequence, and structure-level parameters. For RF-classifiers initially trained on more than 30 parameters, the least predictive parameter is removed until the 30 most-predictive parameters are identified. **(b)** “One vs Rest” receiver operating characteristic curves and precision-recall curves are calculated from the averages of 100 RF-classifiers trained on the 3 different sets of parameters. The shaded region represents 1 standard deviation. **(c)** The importance of individual features for “3D” RF-classifiers is determined. MUT- and MUT+ reflect pre- and post-mutation sequence properties, respectively. The radius of the 3D environment is described where applicable. The interoperability of multiple datatypes in QPSACE allows for the calculation of a mutation’s proximity to the nearest active site—the third most important feature in determining mutant severity. **(d)** QSPACE can be used to analyze mutations found in proteins containing various functional domains. **(e)** The weighted (by phenotype class) area under the curve (AUC) is calculated from the ROC and P-R curves in Panel B for RF-classifiers (3 parameter sets x 100 RF-classifiers) for each functional domain. **(f)** The cumulative importance of residue, sequence, and structure features in “3D” RF-classifiers of mutations in proteins containing each domain.

**Figure 5:**
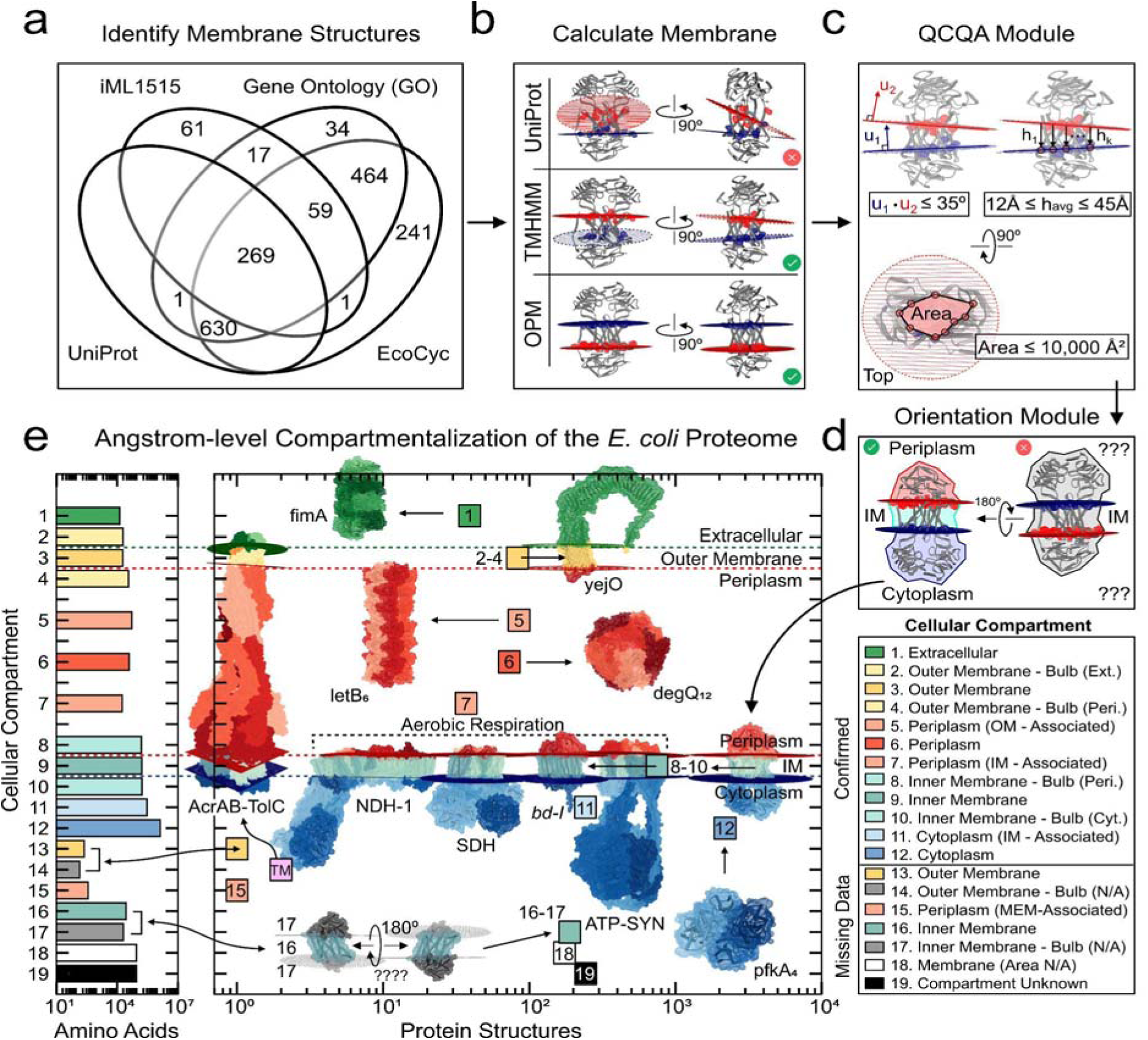
The membrane module orients proteins across the membrane to identify residue-level subcellular compartments of the *E. coli* proteome. **(a)** In the metadata provided by EcoCyc (cellular compartment), Gene Ontology Terms (pathway, function, compartment), iML1515 (metabolic subsystem), and UniProt (topological & transmembrane domains) databases, there are 1,777 protein structures mapped to at least one gene that is associated with the *E. coli* membrane. **(b)** Membrane-crossing residues are identified by the amino acid sequence information provided by UniProt, predicted by DeepTMHMM, and calculated by OPM. From these residues, a plane of best fit is calculated. **(c)** Structures with two calculated membrane planes pass the QCQA analysis if i) the angle between the planes is less than 35°, ii) the thickness of the membrane embedded region is between 12 and 45 Angstroms, and iii) the cross-sectional area of the membrane embedded region is less than 10,000 Å^2^. **(d)** Membrane proteins are oriented using the topological information provided by UniProt (if available) or manually using common protein motifs (see Dataset S6-S7) such that **(e)** the subcellular compartment of every amino acid of the *E. coli* proteome can be determined.

**Figure 6:**
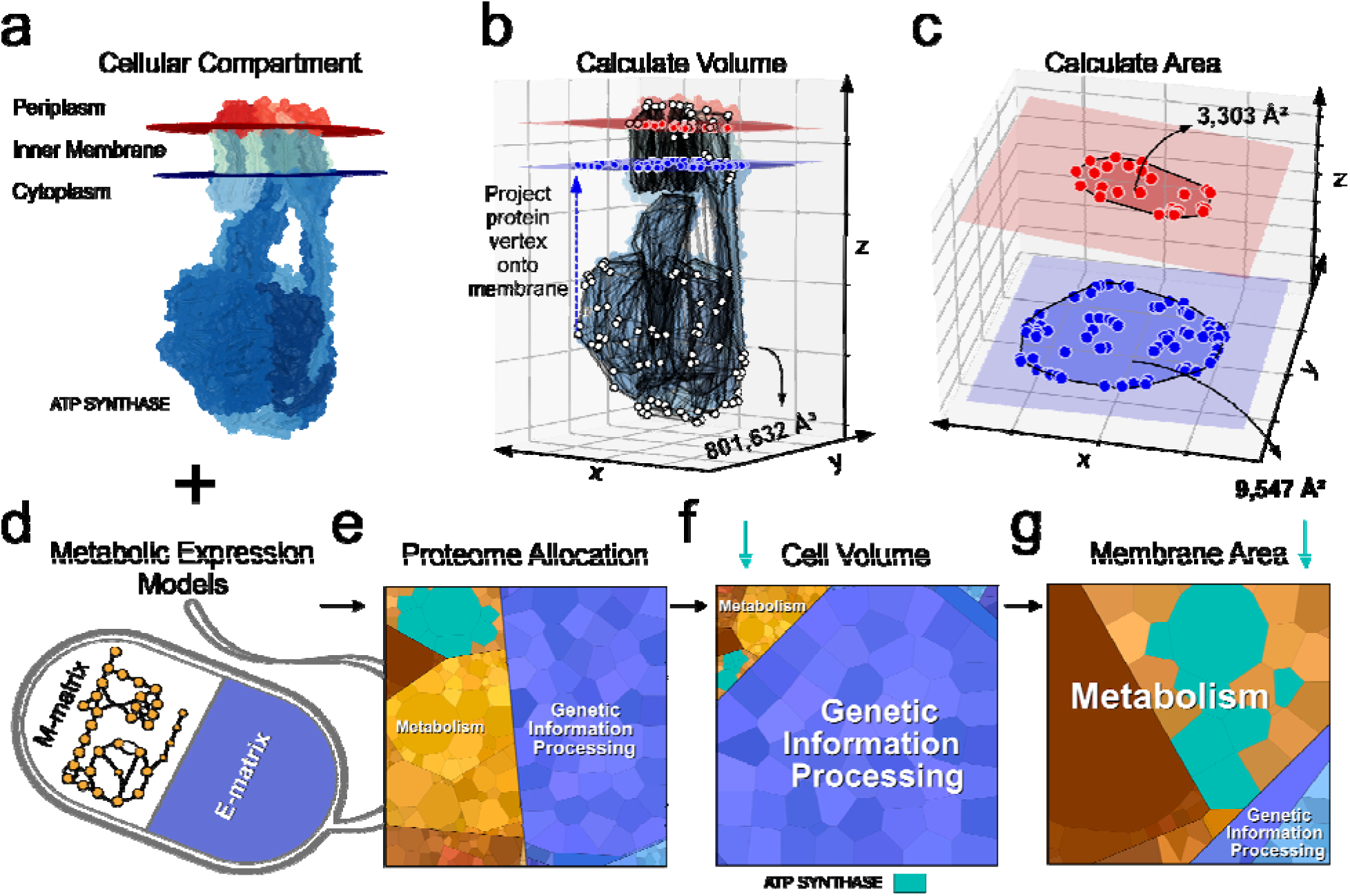
QSPACE integrates with genome-scale models to predict the physical space required by the *E. coli* proteome at optimal growth rate. **(a)** The compartmentalization of each amino acid of the *E. coli* proteome allows for the calculation of geometric properties of all proteins. **(b)** The volume of ATP-synthase and **(c)** its cross-sectional membrane area are shown. **(d)** The integration of QSPACE with genome-scale models (iJL1678b-ME, in this case) of metabolism (M-matrix) and macromolecular expression (E-matrix) (ME-models) can be used to calculate **(e)** the proteome allocation, **(f)** the volumetric allocation, and **(g)** the membrane composition of *E. coli* at optimal growth rate. The expression, volume, and membrane area allocated to ATP-synthase is shown (Panels E-G, cyan). The calculated spatial allocation (Panels F-G) of the macromolecular expression predicted by existing ME-models (in Panel E) is a fundamental advancement towards building genome-scale biophysical whole-cell models.

As an example, we apply QSPACE to the genome of *E. coli* K-12 MG1655 (Fig. 1e, Dataset S1) and identify the quaternary structural representation of its oligomeric proteome—as defined by the multi-decade bibliomic curation available in EcoCyc^39^ and in the *E. coli* genome-scale model iJL1678b^26^. These gene-stoichiometric inputs to the *E. coli* QSPACE are provided in Dataset S2A. Selecting from both experimentally resolved structures deposited in the Protein DataBank (RCSB-PDB)^13^ and from structural models calculated using protein modeling methods (ITASSER^8^, SWISS^10^, AlphaFold^12^ & AlphaFold Multimer^14^) (details in Fig. 2), QSPACE can map the 3D position of 94% (on average) of the amino acids belonging to 3,985 annotated *E. coli* proteins (Fig. 1f, Dataset S3A). The set of structures that QSPACE maps to the *E. coli* structural proteome can be used as 3D scaffolds to map multiple structural, functional, mutational, and spatial data types (Fig. 1g).

### Structural representation of multi-subunit proteins

Proteins often require oligomerization to function properly. The fundamental advancement of the *E. coli* QSPACE over existing genome-scale models with protein structures (e.g. iML1515-GP^40^ see Fig. S1) is that it can be used to identify structures that represent the quaternary shapes of multi-subunit proteins.

To ensure that the user-defined multi-subunit proteins are accurately reflected in the structural data, we designed a pipeline to identify the best available protein structure for a target oligomeric protein, to suggest changes to the user-defined gene stoichiometry when the existing structural data suggests oligomerization, and to generate *de novo* structural models for oligomeric enzymes whose subunits cannot be fully represented by the structures in the PDB. A simplified representation is shown in Figure 2a.

The input to the QSPACE pipeline is a user-defined dictionary of protein complexes and their associated gene-stoichiometries. For *E. coli*, this information is the result of multi-decade bibliomic evidence that has been annotated in the EcoCyc database^39^ and in the genome-scale model iJL1678b-ME^26^ (Dataset S2A). Across these resources of annotated protein-complexes, 31% (1,334/4,309) of *E. coli* genes participate in 1,047 oligomeric complexes, 667 genes are annotated as monomers, and 2,308 genes are not included (i.e. assumed to be monomers) (Fig. S9A-B). In the set of annotated or assumed monomers, QSPACE identified structures (in the PDB or SWISS-MODEL repository) containing one or more oligomeric conformations for 983 of these genes (Fig. 2a.ii & Fig. S9C). QPACE uses a semi-automated pipeline that relies on various structure-derived quality metrics to assess the accuracy of PDB and SWISS-MODELs before redefining the existing monomeric annotation for these genes (see Methods 3.1).

The accuracy of quaternary structures (experimental and modelled) has been the focus of many community-wide structural biology efforts. Previous studies have estimated that the accuracy of the quaternary structures in the PDB (‘biological assemblies’) is in the range of 80-90% ^41–44^, and the accuracy of PISA-generated homo-oligomers to be 85%^45^. QSPACE uses PDB biological assemblies that are author-defined, software-defined (by PISA), or both (see Dataset S8). In cases where the PDB structures suggested oligomerization (contrary to the existing monomeric annotation), we reviewed the publication(s) associated with each PDB structure to confirm the oligomeric structure is believed (by the authors) to be biologically relevant (case IV-V in Fig. S9C-D).

Since 2017, QSQE-scores have been used to assess the quality of oligomeric SWISS-MODELs^46^. Recently, the SWISS-MODEL QSQE-score was shown to distinguish between biologically relevant and non-relevant homodimer structures at a rate of 0.79^15^. Although other modelling platforms perform slightly better^15^, SWISS-MODELs are precomputed and readily available, making them a convenient choice for rapid integration into the QSPACE annotation platform. Thus, in cases where SWISS-MODELs provided structural evidence of oligomerization (cases I-III in Fig. S9C-D), QSPACE relies on the established metrics and thresholds (QSQE^46^ > 0.5, GMQE^47^ > 0.5, and QMN4^47^ > -4) to assess the accuracy of each oligomeric SWISS-MODEL. SWISS-MODELs with scores exceeding these thresholds are used to redefine the oligomerization state of the user-defined monomers.

All oligomeric structures that were considered for changing the annotated *E. coli* monomers are provided in Dataset S2B. The relevant quality metrics associated with each structure ultimately selected in the *E. coli* QSPACE are provided in Dataset S3B.

When structures (in the PDB and/or SWISS-MODEL) are unable to fully reflect the gene-stoichiometry of a user-defined oligomer, the QSPACE platform relies on Alphafold Multimer^14^ (v2.0, via ColabFold^38^) to generate *de novo* structures for desired protein oligomers (Fig. 2.a.iii). Alphafold Multimer (v2.0) was shown to outperform existing methods in modelling physiological homodimers^15^ and has been reported to generate high-confidence homo-oligomeric structures for various organisms, including *E. coli*^16^. QSPACE assigns confidence to AF-Multimer models using an established scoring metric^14^ (0.8*iPTM + 0.2*PTM ≥ 0.8) (Fig. S11). It is important to note that the iPTM thresholds were shown to correlate with biologically relevant homo-dimer models^15^.

Furthermore, we confirmed the physiological relevance of 86% (841/973) of the homo-oligomeric structures that QSPACE ultimately selects to represent *E. coli* proteome (Fig. S10) using QSalignWeb^45^ — a webserver that uses superposition of structures to infer the physiological relevance of a quaternary structure. We provide all relevant quality metrics associated with each structure, and the QSalign inferred relevance (when applicable) for all proteins in the *E. coli* QSPACE in Dataset S3B.

The final structural representation of the *E. coli* proteome is a collection of experimental structures (deposited in the PDB) and models (generated by SWISS-MODEL, I-TASSER, AlphaFold, and AlphaFold Multimer) (Fig. 2b). The collection of structures identified by QSPACE captures the multi-subunit assembly of 1,473 oligomeric proteins (Fig. 2c). Proteins that are not known to oligomerize and that have no structural evidence of oligomerization are mapped to their respective monomeric structures as in previous GEM-PRO formulations^27–31^. We show that QSPACE identifies the structures of higher-order oligomeric enzymes (Fig. 2d). Among these oligomers, the QSPACE platform identifies high-confidence structures for 51/54 ATP-binding cassette (ABC) transporters in *E. coli* (defined in EcoCyc^39^ and/or iJL1678b^26^). Only 4 of these transporters have experimentally resolved structures in the PDB (2QI9, 3RLF, 7CGE & GMHU). We present high-confidence novel structures QSPACE generated with AF-multimer for the 47/50 remaining ABC-transporters in Figure 2e. Incomplete AF-multimer models (3/50, *asterisks* in Fig. 2e) provide obvious suggestions for the correct gene-stoichiometry of ABC-transporters that were incorrectly annotated at the time of publication (e.g. putative ABC-55 transporter is missing an ATP-binding subunit).

When compared to the latest *E. coli* genome-scale model with protein structures (iML1515-GP^40^), QSPACE improves the oligomeric structural annotation for 70% of genes in iML1515, while offering a 2.86-fold increase in gene coverage and higher quality structures (Fig. S1). To our knowledge this result is the most advanced genome-scale structural representation of the *E. coli* proteome and *de facto* represents a major advancement in genome annotation.

### Interoperable data types form the basis for predicting mutant phenotypes

An accurate 3D structural representation of the proteome can serve as a scaffold for mapping multiple data types, thus providing a structured approach to data integration. The interoperability of multiple datatypes can accelerate our understanding of structure-function relationships and mechanisms. To this end, QSPACE uses third-party software (Table S1) to map residue-level, sequence-level, and protein-level properties (columns in Dataset S1) to all amino acids of the *E. coli* proteome. To illustrate the extensive functional content contained in the *E. coli* QSPACE, we provide a global accounting of all functionally important regions of the *E. coli* proteome (Fig. S4 and Dataset S10).

Non-synonymous mutation—the swapping of one amino acid residue for another—provides an opportunity for QSPACE to be used for mutant analysis. Residue-level properties (e.g., the Grantham score^48^) of each mutation are calculated. The physio-chemical properties (e.g., the hydrophobicity) of the local sequence (i.e., 5 amino acids centered at the mutation) of each mutation are also determined. Using the protein structure, QSPACE can calculate the properties of the local 3D environment (all amino acids within a fixed radius) of a mutation. The interoperable mapping of multiple datatypes onto the protein structure also allows for the calculation of unique properties (e.g., the distance between a mutation and the nearest protein active site). A graphical summary of mutant-specific properties can be found in Figure 3a.

The UniProt knowledgebase^37^ contains annotated phenotypes for over 12,000 non-synonymous *E. coli* mutations. We use keyword phrases (Dataset S4) to assign each mutations annotated phenotype to one of eight phenotype classes of varying severity (Fig. 3b). Combined with the 100 mutant-specific properties calculated by QSPACE (Dataset S5), the mutant-phenotype UniProt dataset can be used to train Random Forest (RF) classifiers that can predict the severity of mutations in novel mutational databases (e.g., from adaptive laboratory evolutions, the long-term evolution experiment, or the natural sequence variants) (Fig. 3c).

### Random-forest classification of mutant phenotypes

We investigated the accuracy with which Random Forest (RF) classifiers predicted mutant phenotypes and the relative importance of higher-dimensional features. To this end, we selected all UniProt mutations found in proteins containing annotated *active sites* (e.g., Fig. 4a) and calculated 100 mutant-specific properties for each mutation (see Fig. 3, Dataset S5). To quantify the importance of higher-dimensional properties, we trained three sets of RF-classifiers on varying combinations of residue (“0D”, gray), sequence (“1D”, yellow), and structure (“3D”, green) features, and iteratively removed the least-predictive feature after 100 train/test cycles until the 30 most-predictive features were identified (Fig. 4a). For each set of RF-classifiers, we quantified model performance (accuracy, precision, and recall) using “One vs Rest” validation for each phenotype class (Fig. 4b). The importance of individual features used to train the “3D” RF-classifiers is shown (Fig. 4c).

Mutations in the UniProt dataset are not limited to proteins containing active sites (Fig. 4d). Thus, we followed the procedure described in Figure 4a-c to obtain a global assessment of our ability to predict mutant phenotypes found in proteins containing various functionally important annotations (UniProt “Feature”). The Area Under the Curve (AUC) for the Receiver Operating Characteristic (ROC) and Precision-Recall (P-R) curves were weighted by the relative occurrence of each phenotype class (i.e., horizontal line in Figure 4b, right) and plotted for RF-classifiers trained for each functional class (Fig. 4e). For each functional annotation, the cumulative importance of residue-level, sequence-level, and structural features in “3D” RF-classifiers supports the use of protein structures as a context to study interoperable datatypes and mutations (Fig. 4f).

### The membrane module yields angstrom-level subcellular compartmentalization of the *E. coli* proteome

While mapping data types to individual protein complex structures can prove useful, understanding the location and space that these protein complexes occupy in the cell is important for building a genome-scale representation that reflects the physical embodiment of a proteome. To date, genomic databases of *E. coli* (e.g., EcoCyc) assign the entire gene to a subcellular compartment. UniProt sometimes offers sequence annotation of transmembrane and topological (‘bulb’) domains, however, these annotations may be inaccurate (see Fig. 5b) or missing entirely. Since the structure is not used to determine a protein’s subcellular compartment, assigned cellular compartments can often be incomplete (e.g. there is no distinction between membrane proteins that contain and those that do not contain membrane-spanning regions) and the residue-level orientation of a protein across the cell membrane cannot be achieved. Likewise, sequence-based prediction software (e.g., DeepTMHMM^49^) and structure-based prediction software (e.g., OPM^50^) are agnostic to membrane orientation and can also generate erroneous results.

To achieve a residue-level representation of the *E. coli* proteome, we use a structure-guided approach that combines and assesses all available annotations and predictions (from UniProt, DeepTMHMM, and OPM) to better identify the integration and orientation of the membrane-embedded proteome.

QSPACE queries the available gene-level subcellular compartment information provided by Ecocyc^39^, UniProt^37^, Gene Ontology^51^, and genome-scale model iML1515^40^ to identify all potential membrane-embedded protein structures (Fig. 5a). For each identified structure, QSPACE determines the membrane-spanning residues for each subunit using the sequence annotations provided in UniProt, the sequence-based predictions generated by DeepTMHMM^49^, and the structure-based calculation of the membrane planes predicted by OPM^50^. For each of the three sources of residue information (when available), QSPACE calculates the normal vectors of the corresponding membrane planes (Fig. 5b). For each pair of membrane planes, the angle between the planes, and the thickness and area of the membrane-embedded region are used to determine whether the calculated membranes are viable (Fig. 5c).

QSPACE segregates each viable membrane protein into three sections: a membrane-embedded region and two ‘bulbous’ regions. Each bulb is automatically assigned (Dataset S6) to either the cytoplasmic, periplasmic, or extracellular side of either the inner or outer membrane, using the annotated topological domains in UniProt or manually assigned (Dataset S7) using common 3D motifs in the protein structures (Fig. 5d). Proteins annotated to the cell membrane (Fig. 5a) that do not contain a membrane-embedded region are considered ‘membrane-associated’ and tagged to their respective membrane while those tagged to the cytoplasm or periplasm are left unchanged. The gene ontology (GO) terms of genes mapped to non-membrane proteins were used to assign proteins to the cytoplasm, periplasm, or extracellular space.

In *E. coli*, QSPACE was able to assign 86% of proteins (89% of AAs) to one of twelve subcellular compartments (Fig. 5e), resulting in a residue-level annotation of cellular compartmentalization of the *E. coli* proteome across both cellular membranes. The membrane integration for an additional 5% of proteins (2% of AAs) is known (Fig. 5e, compartments #13-17), however there is insufficient information to properly orient these proteins across the membrane (e.g. short, single-pass transmembrane helix proteins, see Dataset S7). Incorporated into genome-scale models that compute protein expression (or proteomic datasets), the residue-level compartmentalization of each protein structure provides a first-principles approach to compute the location and size taken up by a cell’s proteome.

### Computing the physical space required by the *E. coli* proteome

The multi-subunit protein complexes carry out metabolic reactions, transport nutrients across the cell membrane, maintain cellular homeostasis, replicate the cellular genome, and even synthesize other proteins. Considering all these functions simultaneously calls for the use of computational models. As genome-scale models (GEMs) have increased in scope and mechanistic detail^22–26,52^, they require the biosynthesis and proper assembly of multi-subunit complexes to drive the reactions in their reconstructed metabolic networks.

While genome-scale models using protein structures (GEM-PROs) have been used for a variety of applications^53^ (e.g., contextualization of disease-associated human mutations^32^, identification of protein-fold conservation in similar metabolic reactions^31^, prediction of thermosensitivity in a metabolic network^30^, comparative structural analyses of multiple organisms^28^), the promise of a complete physical representation of a functioning cellular proteome has yet to be delivered. QSPACE moves us close to this goal by calculating the subcellular compartment of every amino acid across the proteome.

The successful annotation and 3D orientation of proteins across the subcellular compartments is crucial for building genome-scale models that can predict the physical distribution of the cellular proteome. A geometric analysis of the compartmentalized proteome (Fig. 6a) allows us to calculate the volume occupied (Fig. 6b) as well as the membrane area required (if applicable) (Fig. 6c) by each protein. Genome-scale models of metabolism and macromolecular expression (ME-models) (Fig. 6d) predict the proteome allocation required to sustain growth in optimally growing bacterial cells (Fig. 6e). In calculating the physical space required by each protein, the spatial requirements of model-predicted proteomes can also be determined (Fig. 6f-g). Thus, it is now possible to compute the composition and location of the structural proteome. A more detailed supra-protein-complex-level 3D arrangement requires additional considerations^54–55^.

## Discussion

The 3D visualization and modeling of the structural proteome of a functioning cell has been an implicit goal of genome-scale annotations and computational biology methods. QSPACE, introduced here, rapidly identifies and annotates multi-subunit protein structures (including *de novo* annotations of protein-complex assemblies and *de novo* structural models) at the genome-scale for computational modeling and structural analyses. In conjunction with mutational databases, functional annotations, and other data types, the oligomeric structures identified through QSPACE can be used to obtain a deeper understanding of whole-cell functions.

To achieve a physical representation of the cellular proteome, the structure of each individual protein complex in its native state is needed. To this end, QSPACE allows for multi-gene mapping to oligomeric crystallographic depositions (e.g., PDB bioassemblies), existing homo-oligomeric structural models (e.g., high-quality^47^ SWISS-PROT models^10^ with high QS-scores^46^), and *de novo* high quality oligomeric models (from Alphafold Multimer^14^/ColabFold^38^). Unlike a purely annotative workflow, QSPACE uses a structure-guided assessment to identify previously unannotated oligomeric assemblies and generate *de novo* structural models when the existing structural data for a protein complex is incomplete. As an example, we present the novel structures of 50/54 ABC-transporters in *E.coli*, and show that even incomplete models 3/50 can provide clues to the correct oligomerization of a protein (Fig. 2e). In the *E. coli* QSPACE, we confirmed the physiological relevance for 86% of the homo-oligomeric structures with QSalignWeb^45^ (Fig. S10). Thus, QSPACE achieves the structural representation of multi-subunit protein complexes, a significant advancement over existing genome-scale models using structural biology software (ssbio^27^, see Fig. S1).

The protein structures identified by QSPACE are a well-suited 3D scaffold on which to calculate protein properties, identify enzymatic domains, and analyze impactful mutations. QSPACE’s interoperability of various data types (columns in Dataset S1), can drive biological discovery. In this study, we showed how the residue-level, sequence-level, and protein-level properties calculated by QSPACE (Fig. 3a) for the mutations annotated in the UniProt knowledgebase (Fig. 3b) can be used to accurately predict mutant phenotypes (Fig. 4b & 4e). Interestingly, when we iteratively removed the least predictive properties from the RF-classifiers during the training phase (Fig. 4a), we found that the predictive power of RF models was overwhelmingly the result of structure-level features (Fig. 4c & 4f). Thus, QSPACE provides users a rapid way to interact with relevant structures and interoperable datatypes to elucidate structure-function relationships across multiple scales.

In addition to the annotated mutations in UniProt, QSPACE can also be used to analyze novel mutational data sets from adaptive laboratory evolutions (ALEdb^20^, Fig. S5), the long-term evolution experiment (LTEE^33^, Fig. S6), and the natural sequence variation^36^ of *E. coli* in three dimensions (Fig. 3c). To our knowledge, the *E. coli* QSPACE provides the first 3D representation of the natural sequence variation of an organism at the genome-scale, and it moves the description and scale of the structural proteome to the species level.

QSPACE advances whole cell modeling efforts^54,56–57^ by establishing structural annotations relevant for molecular processes. Advancements with computational genome-scale models (GEMs) over the past decade have allowed for the prediction of proteome allocation for cells at optimal growth rate^22–26,52^. Increasingly detailed, GEMs include reactions for protein assembly and translocation across subcellular compartments (e.g., membranes), however, previous GEM formulations with monomeric protein structures (GEM-PROs) have yet to reflect the biophysical embodiment of these *in silico* processes.

Using a structures-based approach that combines and assesses all available annotations and predictions for membrane-spanning proteins, QSPACE determines the membrane integration and orientation of proteins across both the inner and outer membrane of *E. coli*. In fact, QSPACE even calculates membrane integration for proteins that span both membranes (e.g., AcrAB-TolC efflux pump, PDB:5v5s, see Fig. 5e). In this study, QSPACE determined the subcellular compartment for 89% of amino acids in *E. coli*. As a proof-of-concept, we combine the protein-level information in QSPACE with a genome-scale model of macromolecular expression (iJL1678b-ME^26^) to calculate the physical size occupied by the predicted proteome of *E. coli* at optimal growth rate. To our knowledge, this first-principles approach resulted in the first GEM-PRO that embodies the spatial allocation of the *E. coli* proteome.

Taken together, the QSPACE genome annotation platform proves users a rapid method to interact with the best available quaternary structures for any list of proteins (e.g., a strain), can accommodate natural sequence variations described by the alleleome^36^ to generate species-level structural proteomes, and enables a physical embodiment of the structural proteome against the 3D morphology of the bacterial cell. The analysis of mutant phenotypes and the size calculation for the *E. coli* proteome demonstrate that QSPACE is amenable to diverse applications. As structures are resolved for large protein complexes, as the scope of genome-scale models expands to include an increasing number of niche cellular mechanisms (e.g., stress responses), and as new mutations of functional importance are annotated in publicly available databases, the QSPACE platform will provide an interoperable pipeline for the structural proteomes for a growing list of organisms.

## Limitations

We emphasize that QSPACE is a large-scale annotation platform that interfaces with numerous third-party software to quickly map multiple interoperable datasets onto relevant protein structures. QSPACE applications range from the single-protein to the genome-scale. As such, the structures identified by QSPACE reflect the gene-stoichiometry of the protein complexes defined by the user. For extensively studied organisms (e.g., *E. coli*), these protein complexes have been defined over decades of published work. For less-studied organisms, a structural proteome assigned by the workflow presented in this study may be incomplete. In such cases, QSPACE can still provide insights into the structural proteome. For instance, QSPACE can generate the entirety of all homo-oligomerization states of an organism’s genome. By modifying the user-defined protein complexes to reflect the gene-stoichiometry of any theoretical homo-oligomerization state of a gene(s), QSPACE will identify (or use AF-multimer to generate) structures for these oligomers, provide the user with a quality assessment of each oligomeric structure, and would thus reveal homo-oligomerization states backed by structural evidence.

As QSPACE relies on third-party software and repositories for the generation of novel structures, the mapping of datasets, and the calculation of structural properties, it is limited by the maintenance, capabilities and accuracy of such resources. For example, the use of AF-multimer via ColabFold allows for modelling protein complexes up to 2000 amino acids, and the quality assessment of such models is currently based on their iPTM and PTM scores, rendering QSPACE incapable of generating higher-order structures that have not been published in repositories. As new modelling platforms, better scoring methods, and larger repositories of pre-computed structures are disseminated by the structural biology community, we see potential for their incorporation into the QSPACE workflow to identify increasingly accurate structures for user-defined proteins. Thus, the maintenance of the QSPACE codebase is vital to ensure that QSPACE can provide users with the most-accurate protein structures for future applications.

## Acknowledgements

We would like to thank Marc Abrams for assistance with manuscript editing. This work was funded by Novo Nordisk Foundation (Grant Number NNF20CC0035580) (E.A.C. and B.O.P.) and NIH (Grant R01 GM057089) (B.O.P.).

## Author Contributions

E.A.C. and B.O.P. designed and performed the research; E.A.C., N.M. and M.L. contributed analytical tools and code; E.A.C. and B.O.P. analyzed data; E.A.C. and B.O.P. wrote the manuscript; E.A.C., N.M., M.L. and B.O.P. edited the manuscript; and B.O.P. supervised the research.

## Materials and Methods

Detailed methods are provided in the Supplementary Appendix. Additional information can be found at github.com/EdwardCatoiu/QSPACE.

### Overview of the QSPACE workflow

We encourage the reader to familiarize themselves with the ‘demo_QSPACE.ipynb’ tutorial notebook available at (https://github.com/EdwardCatoiu/QSPACE). The QSPACE platform offers the user flexibility in the use of some or all structural repositories (for identifying/generating structures), third-party software (for calculating structural properties), and mutant datasets. This section will describe the generation of the *E. coli* QSPACE (Dataset S1) and the two applications presented in this study.

The overall QSPACE workflow can be summarized: 1) A list of 4,309 genes identified across 2,661 *E. coli* strains^36^ and the gene-stoichiometries of *E. coli* proteins are annotated in EcoCyc^39^ and iJL1678b^26^ and serve as the user-defined input to the *E. coli* QSPACE; 2) UniProt IDs were identified and corresponding .fasta and .txt files were downloaded; 3) Homology models corresponding to the UniProt IDs are downloaded from various repositories; 4) PDB structures (and associated PDB bioassemblies) corresponding to the UniProt IDs and sequences were downloaded using PDB APIs; 5) For proteins that are annotated to be monomers and for those not included in EcoCyc and/or iJL1678b (assumed monomers), a semi-automated module is used to assess if the oligomeric structures (from PDB/SWISS-MODEL) provide overwhelming evidence of oligomerization; 6) For annotated oligomers (user-defined in #1) whose gene-stoichiometry is not reflected in PDB or SWISS-MODEL, AF-multimer v2.0 (via ColabFold) is used to generate oligomeric models; 7) iPTM and PTM scores are used to assess the quality of AF-multimer models; 8) The highest sequence identity structure (after structural QCQA relevant to its source) is selected for each unique gene stoichiometry. When multiple structures provide the same sequence identity for the same gene stoichiometry, preference is given to PDB, AF-Multimer, AlphaFoldDB, SWISS, and ITASSER, respectively); 9) A structure (or combination of structures) is selected as the representation of each user-defined (or re-defined in #5) protein complex; 10) A CSV file (the backbone of Dataset S1, ‘the QSPACE’) is generated where each row provides a mapping between each amino acid across 4,309 *E. coli* genes (user-defined in #1) and its 3D position (chain and residue number) on the protein structure that reflects the protein complex (user-defined in #1 and/or redefined in #5); 11) When possible, third-party software is used to identify membrane-embedded amino acids in proteins that are believed to be in (or associated with) the membrane; 12) Proteins with membrane-embedded regions are oriented across the membrane using available topological information (automatically) and/or by manual inspection of common motifs with known orientation; 13) Multiple third-party software is used to calculate protein properties; 14) External mutation databases and functional annotations (in UniProt) are mapped to the QSPACE CSV file (Dataset 1).

Notably, in steps 1-8, the highest quality structure for each unique structure-gene stoichiometry is selected from each resource of homology and experimental structures. In step 9, QSPACE selects the specific structure that best represents each user-defined gene-stoichiometries. All relevant quality metrics associated with all structures in the structure pool are defined in Dataset S9. The structures selected by QSPACE (in step 9) from this pool (and all associated quality metrics), are used to generate the residue-level CSV file described in step 10. The associated quality metrics for all selected structures in the final CSV file are provided in Dataset S8.

The QSPACE CSV (Dataset S1, AA-to-structure mapping in step 10, additional data mapped in steps 11-14) contains information that can be utilized for various applications, for example: 15) The severity of UniProt mutants mapped to the QSPACE was determined using a key-word search of their annotated phenotypes; 16) the AA-level mapping in QSPACE was used to calculate the aggregate properties of the local environment (i.e. the amino acids directly adjacent in sequence and in 3D space) of each mutation; 17) annotated mutant phenotypes and calculated properties were used to train RF-classifiers to predict mutant severity; 18) (Separately) the area taken up by each protein in the membrane was calculated from the membrane-embedded regions and inferred membrane planes generated in step 11; 19) The volume of each protein was calculated; 20) The geometry of all proteins were incorporated into a genome-scale model of macromolecular expression, iJL1678b-ME, to compute the physical space taken up by the model-predicted proteome of *E. coli*.

## Data Availability

All data is freely available from public sources.

Structures selected from the PDB were last downloaded March 5^th^, 2023. We show experimental structures from the PDB with accession numbers 2GRX, 5V5S, 7NYU, 1NEK, 6OQS, 6C53, 1PFK, 6V0C. Structures selected from the SWISS-MODEL *E. coli* repository were last downloaded December 20^th^, 2022. We show SWISS-MODELs with UniProt IDs <u>P33232</u> and <u>P39099</u>. Structures selected from the ITASSER *E. coli* database repository were last downloaded November 3^rd^, 2022. Structures selected from the Alphafold database were last downloaded January 15^th^, 2023. We show Alphafold models with UniProt IDs <u>P33924</u> and <u>P30143</u>. Structures were last modelled using ColabFold on April 6^th^, 2023. We show Alphafold Multimer/ColabFold models for protein complexes with EcoCyc IDs <u>CYT-D-UBIOX-CPLX</u> and <u>ABC-13-CPLX</u>.

Protein complex gene stoichiometry data for *E. coli* is provided by the Public SmartTable in EcoCyc at https://ecocyc.org/group?id=Biocyc12-4862-3584200844 and by genome-scale model iJL1674b-ME at https://github.com/SBRG/ecolime.

ALE mutation data is available at https://aledb.org/. LTEE mutation data is available at https://barricklab.org/shiny/LTEE-Ecoli/. Both mutation datasets were mapped to the *E. coli* genome by Catoiu *et. al*. 2023^36^.

Data generated in this study is provided in the Supplementary Material and/or at https://github.com/EdwardCatoiu/QSPACE/.

Select data generated in this study that exceeds the size limits of GitHub is available at https://drive.google.com/drive/folders/1OkXnPK2YP3WAk62Mmu1p00z2dKiS1HQN?usp=sharing.

## Code Availability

All source code for QSPACE is provided at https://github.com/EdwardCatoiu/QSPACE/. The best way to build a QSPACE is to follow the detailed instructions in the iPython tutorial notebook (“demo_QSPACE.ipynb”).

QSPACE could not be possible without the following:

Python v.3.7.9 (https://www.python.org/); Ssbio v.0.9.9.8 (https://github.com/SBRG/ssbio); Biopython v.1.81 (https://github.com/biopython/biopython); ScanNet (https://github.com/jertubiana/ScanNet); Nglview v.0.11.9 (https://github.com/nglviewer/nglview); Pandas v.1.1.5 (https://github.com/pandas-dev/pandas); SciPy v.1.5.4 (https://github.com/scipy/scipy); NumPy v.1.19.5 (https://github.com/numpy/numpy); Matplotlib v.3.3.3 (https://github.com/matplotlib/matplotlib); Matplotlib_venn v.0.11.9 (https://github.com/konstantint/matplotlib-venn); PyVenn (https://github.com/tctianchi/pyvenn); Seaborn v.0.10.1 (https://github.com/mwaskom/seaborn); scikit-learn v.1.0.2 (https://scikit-learn.org/stable/)

## References

1. X. Bai, G. McMullan, S.H.W. Scheres, How cryo-EM is revolutionizing structural biology. Trends in Biochem. Sci. 40(1), 49–57 (2015). DOI: 10.1016/j.tibs.2014.10.005.

2. Y. Cheng, Single-particle cryo-EM—How did it get here and where will it go. Science, 361(6405), 876–880 (2018). DOI: 10.1126/science.aat4346.

3. J.P. Renaud, A. Chari, W. Liu, H.W. Remigy, H. Start, C. Wiesmann, Cryo-EM in drug discovery: achievements, limitations and prospects. Nat. Rev. Drug Discov. 17, 471–492 (2018). 10.1038/nrd.2018.77.

4. T. Nakane, A. Kotecha, A. Sente, G. McMullan, S. Masiulis, P.M.G.E. Brown, I.T. Grigoras, L. Malinauskaite, T. Malinauskas, J. Miehling, T. Uchański, L. Yu, D. Karia, E.V. Pechnikova, E. de Jong, J. Keizer, M. Bischoff, J. McCormack, P. Tiemeijer, S.W. Hardwick, D.Y. Chirgadze, G. Murshudov, A.R. Aricescu, S.H.W. Scheres, Single-particle cryo-EM at atomic resolution. Nature. 587, 152–156 (2020). 10.1038/s41586-020-2829-0

5. Y. Cheng, Membrane protein structural biology in the era of single particle cryo-EM. Curr. Opin. Struct. Biol. 52, 58–63 (2018). 10.1016/j.sbi.2018.08.008

6. W. Zheng, C. Zhang, Y. Li, R. Pearce, E.W. Bell, Y. Zhang, Folding non-homology proteins by coupling deep-learning contact maps with I-TASSER assembly simulations. Cell Rep. 1, (2021). 10.1016/j.crmeth.2021.100014

7. J. Yang, R. Yan, A. Roy, D. Xu, J. Poisson, Y. Zhang. The I-TASSER Suite: Protein structure and function prediction. Nat. Methods, 12, 7–8 (2015). 10.1038/nmeth.3213

8. J. Yang, Y. Zhang. I-TASSER server: new development for protein structure and function predictions. Nucleic Acids Res., 43, W174–W181 (2015). 10.1093/nar/gkv342

9. A. Waterhouse, M. Bertoni, S. Bienert, G. Studer, G. Tauriello, R. Gumienny, F.T. Heer, T.A.P de Beer, C. Rempfer, L. Bordoli, R. Lepore, T. Schwede, SWISS-MODEL: homology modeling of protein structures and complexes. Nucleic Acids Res. 46, W296–W303 (2018). 10.1093/nar/gky427

10. S. Bienert, A. Waterhouse, T.A.P. de Beer, G. Tauriello, G. Studer, L. Bordoli, T. Schwede, The SWISS-MODEL Repository - new features and functionality. Nucleic Acids Res. 45, D313–D319 (2017). 10.1093/nar/gkw1132

11. J. Jumper, R. Evans, A. Pritzel, et al. Highly accurate protein structure prediction with AlphaFold. Nature (2021). 10.1038/s41586-021-03819-2

12. M. Varadi, S. Anyango, M. Deshpande, et al. AlphaFold Protein Structure Database: massively expanding the structural coverage of protein-sequence space with high-accuracy models. Nucleic Acids Res. 50(D1), D439–D444 (2022). 10.1093/nar/gkab1061

13. H.M. Berman, J. Westbrook, Z. Feng, G. Gilliland, T.N. Bhat, H. Weissig, I.N. Shindyalov, P.E. Bourne. The Protein Data Bank. Nucleic Acids Res. 28, 235–242 (2000).10.1093/nar/28.1.235

14. R. Evans, M. O’Neill, A. Pritzel, et al., Protein complex prediction with AlphaFold-Multimer. Preprint. 10.1101/2021.10.04.463034

15. H. Schweke, Q. Xu, G. Tauriello, L. Pantolini, T. Schwede, F. Cazals, A. Lhéritier, J. Fernandez-Recio, L.A. Rodríguez-Lumbreras, O. Schueler-Furman, J.K. Varga, B. Jiménez-García, M.F. Réau, A.M.J.J. Bonvin, C. Savojardo, P.L. Martelli, R. Casadio, J. Tubiana, H.J. Wolfson, R. Oliva, D. Barradas-Bautista, T. Ricciardelli, L. Cavallo, C. Venclovas, K. Olechnovič, R. Guerois, J. Andreani, J. Martin, X. Wang, G. Terashi, D. Sarkar, C. Christoffer, T. Aderinwale, J. Verburgt, D. Kihara, A. Marchand, B.E. Correia, R. Duan, L. Qiu, X. Xu, S. Zhang, X. Zou, S. Dey, R.L. Dunbrack, E.D. Levy, S.J. Wodak, Discriminating physiological from non-physiological interfaces in structures of protein complexes: A community-wide study. Proteomics. 23(17) (2023). 10.1002/pmic.202200323.

16. H. Schweke, M. Pacesa, T. Levin, C.A. Goverde, P. Kumar, Y. Duhoo, L.J Dornfeld, B. Dubreuil, S. Georgeon, S. Ovchinnikov, D. N Woolfson, B.E. Correia, S. Dey, E.D. Levy. An atlas of protein homo-oligomerization across domains of life. Cell (2024).

17. H.H. Wang, F. J. Isaacs, P. A. Carr, Z.Z. Sun, G. Xu, C.R. Forest, G.M. Church, Programming cells by multiplex genome engineering and accelerated evolution. Nature, 460(7257), 894–898 (2009). 10.1038/nature08187

18. T.E. Sandberg, M.J. Salazar, L.L. Weng, B.O. Palsson, A.M. Feist, The emergence of adaptive laboratory evolution as an efficient tool for biological discovery and industrial biotechnology. Metab. Eng. 56, 1–16 (2019). 10.1016/j.ymben.2019.08.004.

19. K. Kim, M. Kang, SH Cho, E. Yoo, U. Kim, S. Cho, B. Palsson, BK Cho, Minireview: Engineering evolution to reconfigure phenotypic traits in microbes for biotechnological applications, Comput. Struct. Biotechnol. J. 21, 563–573 (2023). 10.1016/j.csbj.2022.12.042.

20. P.V. Phaneuf, D. Gosting, B.O. Palsson, A.M. Feist, ALEdb 1.0: a database of mutations from adaptive laboratory evolution experimentation. Nucleic Acids Res. 47(D1) D1164–D1171 (2019). 10.1093/nar/gky983

21. J.D. Tibocha-Bonilla, C. Zuñiga, A. Lekbua, C. Lloyd, K. Rychel, K. Short, K. Zengler, Predicting stress response and improved protein overproduction in Bacillus subtilis. NPJ Syst. Biol. Appl. 8, (2022). 10.1038/s41540-022-00259-0

22. E.J. O’Brien, J.A. Lerman, R.L. Chang, D.R. Hyduke, B.Ø. Palsson, Genome-scale models of metabolism and gene expression extend and refine growth phenotype prediction. Mol. Syst. Biol. 9, (2013). https://www.embopress.org/doi/pdf/10.1038/msb.2013.52#sec-18

23. B. Du, L. Yang, C.J. Lloyd, X. Fang, B.O. Palsson, Genome-scale model of metabolism and gene expression provides a multi-scale description of acid stress responses in Escherichia coli. PLOS Comp. Bio. 15(12), (2019). 10.1371/journal.pcbi.1007525

24. K. Chen, Y. Gao, N. Mih, BO Palsson, Thermosensitiviy of growth is determined by chaperone-mediated proteome reallocation. PNAS, 114(43), 11548–11553 (2017). 10.1073/pnas.1705524114

25. L. Yang, N. Nih, A. Anand, J.H. Park, J. Tan, J. Yurkovich, J.M. Monk, C.J. Lloyd, T.E. Sandberg, S.W. Seo, D. Kim, A.V. Sastry, P. Phaneuf, Y. Gao, J.R. Broddrick, K. Chen, D. Heckmann, R. Szubin, Y. Hefner, A.M. Feist, B.O. Palsson, Cellular responses to reactive oxygen species are predicted from molecular mechanisms. PNAS. 116(28), 14368–14373 (2019). 10.1073/pnas.1905039116

26. C.J. Lloyd, A. Ebrahim, L. Yang, Z.A. King, E. Catoiu, E.J. Obrien, J.K. Liu, B.O. Palsson, COBRAme: A computational framework for genome-scale models of metabolism and gene expression. PLOS Comp. Bio, 14(7), (2018). 10.1371/journal.pcbi.1006302

27. N. Mih, E. Brunk, K. Chen, E. Catoiu, A. Sastry, E. Kavvas, J.M. Monk, Z. Zhang, B.O. Palsson, ssbio: a Python framework for structural systems biology. Bioinformatics. 34(12), 2155–2157 (2018). 10.1093/bioinformatics/bty077

28. E. Brunk, N. Mih, J.M. Monk, Z. Zhang, E.J. Obrien, S.E. Bliven, K. Chen, R.L. Chang, P.E. Bourne, B.O. Palsson, Systems biology of the structural proteome. BMC Syst. Biol. 10(26), (2013). 10.1186/s12918-016-0271-6

29. R.L. Chang, L. Xie, P.E. Bourne, B.O. Palsson, Drug off-target effects predicted using structural analysis in the context of a metabolic network model. PLOS Comp. Biol. 6(9), (2010). 10.1371/journal.pcbi.1000938

30. R.L Chang, K. Andrews, D. Kim, Z. Li, A. Godzik, B.O. Palsson, Structural systems biology evaluation of metabolic thermotolerance in Escherichia coli. Science. 34(6137), 1220-1223 (2013). 10.1126/science.1234012

31. Y. Zhang, I. Theile, D. Weekes, Z. Li, L. Jaroszewski, K. Ginalski, A. Deacon, J. Wooley, S. Lesley, I.A. Wilson, B.O. Palsson, A. Osterman, A. Godzik, Three-dimensional structural view of the central metabolic network of Thermotoga maritima. Science, 325(5947), 1544-1549 (2009). 10.1126/science.1174671

32. E. Brunk, S. Sahoo, D.C. Zielinski, A. Altunkaya, A. Dräger, N. Mih, F. Gatto, A. Nilsson, G.A. Preciat-Gonzalez, M.K. Aurich, A. Prlić, A. Sastry, A.D. Danielsdottir, A. Heinken, A. Noronha, P.W. Rose, S.K. Burley, R.M.T. Fleming, J. Nielsen, I. Thiele, B.O. Palsson, Recon3D enables a three-dimensional view of gene variation in human metabolism. Nat. Biotechnol. 36, 272–281 (2018). 10.1038/nbt.4072

33. Barrick Lab, LTEE-Ecoli. [Online]. Available: https://barricklab.org/shiny/LTEE-Ecoli/ [Accessed 1 April (2022].

34. R.E. Lenski, M.R. Rose, S.C. Simpson, S.C. Tadler, Long-term experimental evolution in Escherichia coli adaptation and divergence during 2,000 generations, Am. Nat. 138(6) 1315–1341 (1991).

35. O. Tenaillon, J. E. Barrick, N. Ribeck, D. E. Deatherage, J. L. Blanchard, A. Dasgupta, G.C. Wu, S. Wielgoss, S. Cruveiller, C. Médigue, D. Schneider, R. E. Lenski, Tempo and mode of genome evolution in a 50,000-generation experiment. Nature. 536, 165–170 (2017).

36. E.A. Catoiu, P. Phaneuf, J.M. Monk, B.O. Palsson, Whole genome sequences from wild-type and laboratory evolved strains define the alleleome and establish its hallmarks. PNAS. 120(15), 2023. 10.1073/pnas.221883512

37. R. Apweiler, A. Bairoch, C.H. Wu, W.C Barker, B. Boeckmann, S. Ferro, E. Gasteiger, H. Huang, R. Lopez, M. Magrane, M.J Martin, D.A Natale, C. O’Donovan, N. Redaschi, L.S. Yeh. UniProt: the Universal Protein knowledgebase. Nucleic Acids Res. 32, D115–D119 (2004). 10.1093/nar/gkh131

38. M. Mirdita, K. Schütze, Y. Moriwaki, et al. ColabFold: making protein folding accessible to all. Nat Methods. 19, 679–682 (2022). 10.1038/s41592-022-01488-1

39. I.M. Keseler, J. Collado-Vides, A. Santos-Zavaleta, M. Peralta-Gil, S. Gama-Castro, L. Muñiz-Rascado, C. Bonavides-Martinez, S. Paley, M. Krummenacker, T. Altman, P. Kaipa, A. Spaulding, J. Pacheco, M. Latendresse, C. Fulcher, M. Sarker, A.G. Shearer, A. Mackie, I. Paulsen, R.P. Gunsalus, P.D. Karp, Ecocyc: a comprehensive database of Escherichia coli biology. Nucleic Acids Res. 39, D583–590 (2011). 10.1093/nar/gkq1143

40. J. Monk, C.J. Lloyd, E. Brunk, et al. iML1515, a knowledgebase that computes Escherichia coli traits. Nat Biotechnol. 35, 904–908 (2017). 10.1038/nbt.3956

41. E. Krissinel & K. Henrick, Inference of macromolecular assemblies from crystalline state. J. Mol. Biol. 372, 774–797 (2007). 10.1016/j.jmb.2007.05.022

42. Q. Xu, A.A. Canutescu, G. Wang, M. Shapovalov, Z. Obradovic & R.L. Dunbrack, Statistical analysis of interface similarity in crystals of homologous proteins. J. Mol. Biol. 381, 487–507 (2008). 10.1016/j.jmb.2008.06.002

43. K. Baskaran, J.M. Duarte, N. Biyani, S. Bliven & G. Capitani, A PDB-wide, evolution-based assessment of protein–protein interfaces. BMC Struct. Biol. 14(22) (2014). 10.1186/s12900-014-0022-0

44. E.D. Levy, PiQSi: protein quaternary structure investigation. Structure 15(11), 1364– 1367 (2007). 10.1016/j.str.2007.09.019.

45. S. Dey, J. Priluskiy & E.D. Levy, QSalignWeb: A server to predict and analyze protein quaternary structure. Front Mol Biosci (2022). 10.3389/fmolb.2021.787510

46. M. Bertoni, F. Kiefer, M. Biasini, L. Bordoli, T. Schwede, Modeling protein quaternary structure of homo- and hetero-oligomers beyond binary interactions by homology. Sci. Rep. 7 (2017). 10.1038/s41598-017-09654-8

47. P. Benkert, M. Biasini, T. Schwede, Toward the estimation of the absolute quality of individual protein structure models. Bioinformatics 27, 343–350 (2011). 10.1093/bioinformatics/btq662

48. R. Grantham, Amino acid difference formula to explain protein evolution, Science. 185, 862–864 (1974). doi:10.1126/science.185.4154.862

49. J. Hallgren, K.D. Tsirigos, M.D. Pedersen, J.J.A. Armenteros, P. Marcatili, H. Nielsen, A. Krogh, O. Winther. DeepTMHMM predicts alpha and beta transmembrane proteins using deep neural networks. bioRxiv (2022). doi: 10.1101/2022.04.08.487609

50. M.A Lomize, I.D. Pogozheva, H. Joo, H.I. Mosberg, A.L Lomize, OPM database and PPM web server: resources for positioning of proteins in membranes. Nucleic Acids Res. 40, D370–D376 (2012). 10.1093/nar/gkr703

51. M. Ashburner, C.A. Ball, J. A. Blake, et al. Gene ontology: tool for the unification of biology. The Gene Ontology Consortium. Nat. Genet. 25(1), 25–29 (2000). 10.1038/75556

52. J.K. Liu, E.J. O’Brien, J.A. Lerman, K. Zengler, B.O. Palsson & A.M. Feist, Reconstruction and modeling protein translocation and compartmentalization in Escherichia coli at the genome-scale. BMC Syst. Biol. 8(110), 2014. 10.1186/s12918-014-0110-6

53. N. Mih, B.O. Palsson, Expanding the uses of genome-scale models with protein structures. Mol. Cyst. Biol. 15(11), (2019). 10.15252/msb.20188601

54. Z.R. Thornburg, D.M. Bianchi, T.A. Brier, B.R. Gilbert, T.M. Earnest, M.C.R. Melo, N. Safronova, J.P. Sáenz, A.T. Cook, K.S. Wise, C.A. Hutchison, H.O. Smith, J.I. Glass, Z. Luthey-Schulten, Fundamental behaviors emerge from simulations of a living minimal cell. Cell. 185(2), 345–360, 2022 10.1016/j.cell.2021.12.025.

55. M. Maritan, L. Autin, J. Karr, M.W. Covert, A.J. Olson, D.S. Goodsell, Building Structural Models of a Whole Mycoplasma Cell. J. Mol. Biol. 434(2) 2022. 10.1016/j.jmb.2021.167351.

56. E.J. O’Brien, J.M. Monk, B.O Palsson. Using Genome-scale Models to Predict Biological Capabilities. Cell. 161(5), 971–987 (2015). doi: 10.1016/j.cell.2015.05.019.

57. J.R Karr, J.C. Sanghvi, D.N. Macklin, M.V. Gutschow, J.M. Jacobs, B. Bolival Jr, N. Assad-Garcia, J.I. Glass, M.W. Covert. A whole-cell computational model predicts phenotype from genotype. Cell. 150(2), 389–401 (2012). doi: 10.1016/j.cell.2012.05.044.

58. Y. Rose, J.M. Duarte, R. Lowe, J. Segura, C. Bi, C. Bhikadiya, L. Chen, A.S. Rose, S. Bittrich, S.K. Burley, J.D. Westbrook. RCSB Protein Data Bank: Architectural Advances Towards Integrated Searching and Efficient Access to Macromolecular Structure Data from the PDB Archive, J. Mol. Biol. 433(11), (2021). doi: 10.1016/j.jmb.2020.11.003

59. R. Linding, L.J. Jensen, F. Diella, P. Bork, T.J. Gibson, R.B. Russell. Protein disorder prediction: implications for structural proteomics. Structure. 11(11), 1453–1459 (2003). doi: 10.1016/j.str.2003.10.002.

60. J. Tubiana, D. Schneidman-Duhovny, H.J. Wolfson, ScanNet: an interpretable geometric deep learning model for structure-based protein binding site prediction. Nat Methods. 19, 730–739 (2022). 10.1038/s41592-022-01490-7

61. J. Cheng, A.Z. Randall, M.J. Sweredoski, P. Baldi, SCRATCH: a protein structure and structural feature prediction server. Nucleic Acids Res. 33(W), 72–76 (2005). doi: 10.1093/nar/gki396.

62. W. Kabsch and C. Sander, Dictionary of protein secondary structure: Pattern recognition of hydrogen-bonded and geometrical features. Biopolymers, 22, 2577–2637 (1983). 10.1002/bip.360221211

63. M.F. Sanner, A.J. Olson, J.C. Spehner, Reduced Surface: An Efficient Way to Compute Molecular Surfaces. Biopolymers. 38, 305–320 (1996).

64. W. Liebermeister, E. Noor, A. Flamholz, D. Davidi, J. Bernhardt, and R. Milo, Visual account of protein investment in cellular functions. PNAS. 111(23), 8488–8493 (2014). 10.1073/pnas.131481011

